# Animal-to-Animal Variability in Hippocampal Remapping

**DOI:** 10.1101/2020.12.30.424873

**Authors:** Parsa Nilchian, Matthew A. Wilson, Honi Sanders

## Abstract

Hippocampal place cells form a map of an animal’s environment. When the animal moves to a new environment, place field locations and firing rates change, a phenomenon known as remapping. Different animals can have different remapping responses to the same environments. This variability across animals in remapping behavior is not well understood. In this work, we analyzed electrophysiological recordings from Alme et al. (2014), in which five male rats were exposed to 11 different environments. To compare the hippocampal maps in two rooms, we computed average rate map correlation coefficients. We discovered that the heterogeneity in animals’ remapping behavior is structured: animals’ remapping behavior is consistent across a range of independent comparisons. Our findings highlight that remapping behavior between repeated environments depends on animal-specific factors.

## Introduction

Place cells are hippocampal neurons that fire at specific locations in an environment (O’Keefe and Dostrovsky, 1971), leading the place cell population to collectively form a map of the environment (O’Keefe, 1976). When animals are exposed to new environments, the firing patterns of place cells change unpredictably, leading to a new map across the population (Muller and Kubie, 1987). This phenomenon is referred to as ‘‘remapping.” Remapping enables place cell firing patterns to be unique for each environment, thus providing a potential mechanism for encoding novel experiences, and for enabling context-dependent learning (Colgin et al., 2008).

However, the context-identification problem is not as simple as it might seem. Animals don’t have direct access to objective context labels, but instead must *infer* context identity. Characterizing remapping as hidden state inference (Sanders et al., 2020) captures many of the ways in which remapping does not follow “one room = one map” (e.g., Lever et al., 2002; Markus et al., 1995; Law et al., 2016). One particular implication of the animal’s lack of access to objective context labels is that different animals may infer context identity differently when presented with the same experiences. This has been observed previously in studies focusing on other aspects of remapping

**Table 1:** Number of place cells recorded from each animal and animals’ color codes used in the figures of this work.

| Animal ID | Number of place cells recorded | Color code |
| --- | --- | --- |
| 18024 | 25 | red |
| 17769 | 35 | purple |
| 19251 | 38 | orange |
| 17894 | 66 | blue |
| 18237 | 66 | green |

(Lever et al., 2002; Wills et al., 2005) and is widely understood by experimenters, but has not been quantified or directly investigated.

The animal’s lack of access to objective context labels could result in animal-to-animal variability in remapping behavior, as different animals take different approaches to an ambiguous problem. Previous work had shown that factors such as age could cause changes in remapping (Barnes et al., 1997; Lister and Barnes, 2009), and even that variation in remapping among aged animals correlated with other firing phenomena Hok et al. (2012). However, previous work did not investigate the variability in remapping behavior across animals during normal cognitive function.

In this study, we reanalyzed electrophysiological data from Alme et al. (2014), in which rats were exposed to 11 different rooms over 28 sessions. To measure the extent of remapping, we calculated average Rate Map Correlations (RMCs). We verified that RMCs are higher for same-environment comparisons than for different-environment comparisons. However, there was substantial variability in remapping behavior. We found that variability across animals was consistent for a range of comparisons, pointing to consistent individual differences in remapping response. We demonstrate that these individual differences in remapping behavior across animals persist when controlling for several different behavioral parameters and differences in cell yield in the recordings for different animals. Overall, we provide evidence for the hypothesis that individual animals have consistently different remapping responses to the same set of experiences.

## Materials and Methods

The data for this study was collected by Alme et al. (2014), and permission was granted to us to conduct the analyses described below. Detailed descriptions of the animals, surgery, electrode preparation, and implantation can be found in the original publication (Alme et al., 2014). The number of cells recorded from each animal is shown in Table 1.

### Behavioral Procedures

Seven male Long Evans rats foraged for food in one familiar (F) and ten novel (N) environments over a two-day period. 10 of the 11 boxes had dimensions of 100 x 100 x 50 cm, while one box was 100 x 100 x 80 cm. A white cue card was placed on the North wall of each box, but its size and position varied across rooms. Each animal was tested over a period of two days with 8-hour recording sessions on each day. Electrophysiological recordings in each novel room lasted 30 minutes (2 sessions of 15-minutes each), followed by 15-minute rest blocks (see Fig. 1A for schedule of recording day). The familiar room was tested as the first and last session of each day and was preceded and succeeded by a rest period. Five of the ten novel rooms (N1-5) were tested on the first day, and the other five novel rooms (N6-N10) were tested on the second day. The animal was recorded twice in each novel room with no rest period in between. In five of the seven animals, N1 and N6, the first novel rooms of day 1 and 2, were tested at the end of each day again. The data of these five animals were used for this study because we were interested in the repetitions of N1 and N6 in particular. One of the animals was only tested once in N1. The data of animal 17769 room F1 and animal 19251 room N4 session 1 (N4) was missing.

#### Notation of Sessions

The familiar room is denoted as F1 and F1* on day 1 and F2 and F2* on day 2. For the familiar room the number following F indicates the day and the “*” refers to the second session at the end of the day. Novel rooms are described as N followed by the room number (e.g. N3 for room 3) and the immediate repetitions are marked with “!” (e.g. N3!). The third and fourth repetitions of rooms N1 and N6 at the ends of day 1 and 2 are labeled with “*” and “*!” (e.g. N1* and N1*!). In general, “*” denotes sessions in the same room at a different time of the day, whereas “!” denotes sessions that are immediately repeated.

### Data Analysis

We analyzed the data with Python using the Numpy (Harris et al., 2020) and Scipy (Virtanen et al., 2020) libraries. Plots were developed with Matplotlib (Hunter, 2007). The pre-processing steps were based on (Alme et al., 2014).

#### Position Data Processing

The position data consisted of 1D arrays of x- and y-coordinates recorded with an approximate frequency of 25 Hz. X- and y-positions were smoothed with a 1D Gaussian filter (sigma = 5 samples ≈ 200 ms).

#### Behavioral Metrics

To quantify the animals’ behavior in each session, we used the smoothed position data to calculate the mean: 1) speed, 2) acceleration, and 3) absolute angular angular velocity of each animal in each session.

1. **Speed.** The change in x- and y-position (the difference between two consecutive smoothed x- or y-positions respectively) was divided by the time difference between the two position coordinate recordings (approximately 0.04 s). Total speed was the square root of the sum of squares of the x- and y-speed. The speed was then smoothed with a Gaussian filter (sigma = 7 time bins ≈ 0.28 s). Hence, we obtained a 1D speed-vector, with each entry describing the animal’s speed between two consecutive position recordings. This smoothed speed was used for all further analyses.
2. **Acceleration.** The x- and y-speed 1D-vectors were used to calculate the acceleration in x- and y-direction respectively, across all animals and sessions. To calculate the x-acceleration vector, we calculated the difference in two consecutive x-speed entries to obtain the change in speed in the x-direction. Subsequently, we divided the change in speed in the x-direction by the amount of time that passes between the speed entries to obtain a 1D x-acceleration vector. The 1D y-acceleration vector was calculated according to the same steps using the y-speed vector. We then calculated the total acceleration as the square root of the sum of the x-acceleration and y-acceleration squared to obtain a 1D acceleration vector. The animals’ acceleration vector was smoothed with Gaussian filters (sigma = 10 time bins ≈ 0.4 s). Lastly, we computed the mean acceleration for all animals and sessions.
3. **Absolute angular velocity.** The mean absolute angular velocity for each animal in each session was calculated using the session’s smoothed position data. The change in x- and y-position was used to calculate a 1D-vector describing the angle of the animal’s movement between each of the position recording times. Subsequently, we calculated the difference between two consecutive angle entries to obtain a 1D-vector describing the angular change in the direction of the animal across the session. The angular change in direction was divided by the time that had passed between the two position entries to obtain the a 1D-vector describing the animal’s angular velocity across the session. Lastly, we converted each entry of the 1D-angular velocity vector to its absolute value and calculated the mean of the resulting absolute angular velocity vector.

#### Rate Maps

The 100 x 100 cm boxes were divided into 400 (20 x 20) 5×5 cm bins. As is standard in the field, place fields were only calculated using time points when the animal was moving at least 5 cm/s, in order to limit analyses to periods with engaged behavior. For each cell and each spatial bin, the firing rate of that cell was computed as the ratio of the number of spikes fired by that cell when the animal was in that bin (spike count) divided by the total time spent in that bin (occupancy). Both spike counts and occupancy in each bin were calculated using only time points when the smoothed speed (see “Behavioral Metrics” for definition) was above 5 cm/s (fig 2). Rate maps were then smoothed with a 2D Gaussian filter (sigma = 1 bin = 5 cm). This value of sigma was chosen to match the firing rates in figures 3 and S2 in (Alme et al., 2014). Bins with no occupancy were replaced with a gaussian filtered average of surrounding bins with appropriate normalizations. When comparing the place field locations and firing rates of our rate maps to (Alme et al., 2014), we noticed minor differences in the maximum firing rates (commonly deviations of 1-2 Hz), which may have been related to differences in the speed-filtering and smoothing of the data. Overall, we were able to replicate the rate maps of the original publication well (fig 3).

#### Rate Map Correlation (RMC)

We calculated RMCs to quantify the amount of remapping between session pairs. We categorized the cells into three groups for a given session. Cells either had no spikes (silent), less than 10 spikes (below threshold), or at least 11 spikes (active) in a given session. We did not define the RMC if the cell was below threshold or silent in both sessions. The RMC was defined as 0 if the cell was silent in one session and was active in the other. If the cell was active in one session and either active or below threshold in the other, we defined the RMC as follows: we turned the smoothed rate map of each cell in each session into a 1D vector with 400 entries, each corresponding to one of the room’s bins. We then calculated the Pearson Correlation Coefficient for each cell between this 1D vector for the two sessions being compared. This correlation coefficient is the RMC.

#### Average RMC

In addition to calculating the RMC of individual place cells across two sessions, we also averaged across cells for a given session pair to get the average RMC.

#### Animal-to-animal variability

To quantify the animal-to-animal variability in remapping behavior, we performed three General Linear Models (GLMs). We were interested in identifying relationships between the remapping behavior for the repetitions of the familiar room, the repetitions of N1 and N6 at the ends of each day, and the immediate repetitions of the novel rooms (N). We correlated: 1. Mean N between session vs. Mean F between session RMCs, 2. Mean N1 between session vs. Mean N6 between session RMCs, 3. Mean N within session vs. Mean N between session RMCs. Table 2 provides a detailed description of the variables, the respective sessions used, and averaging procedures for this analysis.

**Table 2:** Variables used in GLMs of Figure 7. Each row describes how we calculated the value of a variable used in Figure 7. The rows contain the name of the variable in the left column, a verbal description of the session pairs used in the middle column, and the actual list of session pairs in the right column. For a given variable, we calculated the RMC for each cell for the session pairs listed in the right column. We averaged across all cells for each session pair. We then averaged across all session pairs for each animal. In this way, we arrive at a single value of each variable for each animal.

| Variable | Description | Sessions |
| --- | --- | --- |
| **Mean F between session** | The repetitions of the familiar room at the ends of days 1 and 2. | • F1 / F1*<br>• F1 / F2<br>• F1 / F2*<br>• F1* / F2<br>• F1* / F2*<br>• F2 / F2* |
| **Mean N1 between session** | The repetitions of the novel room N1 at the end of day 1. | • N1 / N1*<br>• N1 / N1*!<br>• N1! / N1*<br>• N1! / N1*! |
| **Mean N6 between session** | The repetitions of the novel room N6 at the end of day 2. | • N6 / N6*<br>• N6 / N6*!<br>• N6! / N6*<br>• N6! / N6*! |
| **Mean N between session** | The repetitions of the of the rooms N1 and N6 at the ends of days 1 and 2 respectively. | • N1 / N1*<br>• N1 / N1*!<br>• N1! / N1*<br>• N1! / N1*!<br>• N6 / N6*<br>• N6 / N6*!<br>• N6! / N6*<br>• N6! / N6*! |
| **Mean N within session** | The immediate repetitions of the novel rooms N1 to N10. | • N1 / N1!<br>• N2 / N2!<br>• N3 / N3!<br>• N4 / N4!<br>• N5 / N5!<br>• N6 / N6!<br>• N7 / N7!<br>• N8 / N8!<br>• N9 / N9!<br>• N10 / N10! |

#### Behavioral Controls

We used the animals’ mean speeds, accelerations, and absolute angular velocities as behavioral parameters to control the relationship between the neural dependent and independent variables of Fig. 7C. The dependent variable in Fig. 7C was the animals’ Mean N within session RMC and the independent variable was the animals’ Mean N between session RMC, both defined in Table 2. To control the findings of Fig. 7C we reconstructed the GLM six times, each time adding a behavioral control parameter to the model as a second independent variable. A detailed description of the behavioral control variables can be found in Table 3. Initially, we used mean differences in mean speeds, accelerations, and absolute angular velocities (Method 1, orange). Subsequently, we used speed means, acceleration means, and absolute angular velocity means, as behavioral independent variables (Method 2, yellow).

## Results

We were interested in looking at the changes in hippocampal maps between experiences. In the experiment we analyzed (Alme et al., 2014), electrophysiological recordings were taken from the hippocampi of rats, while they foraged for food in 11 different 100 cm x 100 cm boxes over two days (Fig 1, see Methods for details). According to the experimental protocol, some rooms were repeated immediately (N1-N10), while others were repeated at a later time during the same day (F, N1 and N6) or during a different day (F). This experiment allowed us to compare repetitions of a variety of different experiences: immediate repetitions of novel rooms, repetitions of novel rooms later during the day, and repetitions of the familiar room. This experiment also allowed us to compare the hippocampal representation between distinct experiences.

First, we had to quantify maps for each experience. The processing pipeline (Fig. 2) illustrates how we combined the cells’ spiking data with the animals’ position data to create rate maps for each place cell across all sessions (see Materials and Methods for more detail).

We observed multiple examples of remapping (Fig. 3). We can use rate map correlations (RMCs) to quantify remapping of individual cells for a particular pair of sessions. High RMCs indicate high similarity in the rate maps of the cell across two sessions and thus less remapping. Low RMCs indicate low similarity in the maps and thus more remapping. In some cases, place fields were conserved but firing frequencies changed, resembling rate remapping (Cell T0505 in sessions N1 and N1*, RMC = 0.69). Global remapping occurred when the place field locations changed across different sessions (T07C02 in animal 19251 between sessions N2 and N3, RMC = −0.14). However, not all place cells remapped across all sessions. In many cases, firing frequencies and place field locations were conserved (T09C06 in animal 17894 sessions N3 and N3!, RMC = 0.82 or T07C02 in animal 19251 between rooms N3 and N3!, RMC = 0.94).

We looked at the RMCs of individual cells across different and repeated sessions to assess the remapping behavior across the place cell population (Fig. 4). For this analysis, we grouped all potential repeated and different session comparisons together. Since it is generally assumed that place cells remap when animals move to new rooms, we expected a distribution of RMCs clustered around 0 for different room comparisons. This pattern was confirmed for all 5 animals, indicating that most place cells do indeed remap across different rooms (Fig. 4). Interestingly, we observed a tendency towards a tail of higher RMCs for the different session comparisons. These relatively high RMCs may correspond to some non-repeated sessions, i.e. comparisons of different rooms, having limited remapping (clusters of high RMCs in Fig. 5). Given that all boxes have the same shape and dimensions these place cells could indicate the similarity of the animals’ context despite the changing rooms. Based on the field’s current understanding of remapping, one would expect only a small number of place cells to remap across repeated sessions (Leutgeb et al., 2005; Colgin et al.,

**Table 3:**
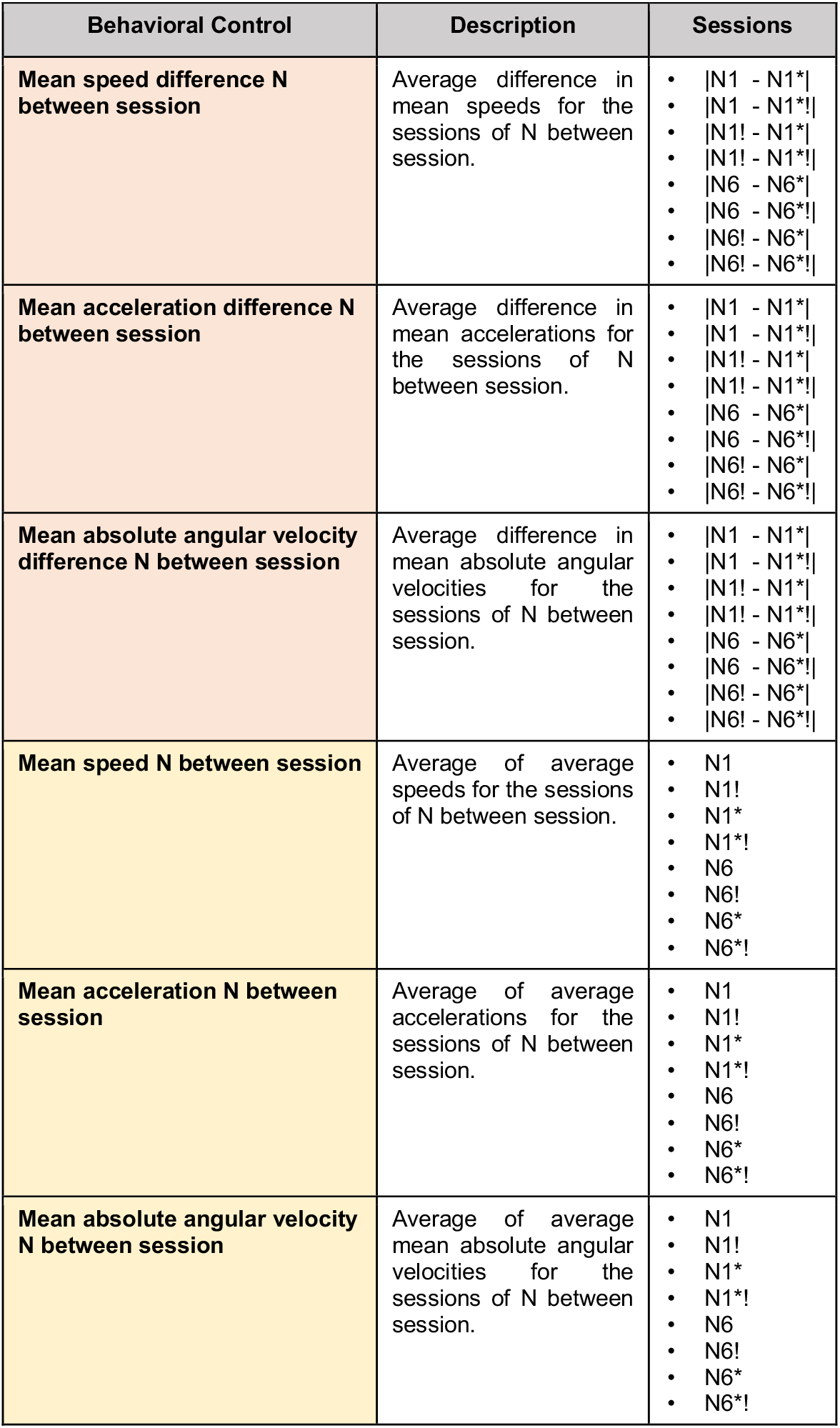
Behavioral controls used for the General Linear Models (GLMs) of Fig. 7C. Each row describes how we calculated the value of a control variable used in Figure 7C. The rows contain the variable’s name in the left column, a verbal description of the variable in the middle column, and the session pairs used in the right column. We controlled the relationship between the dependent and independent variables displayed in 7C by reconstructing the GLM with the addition of a behavioral variable to the model. In total, we used six different behavioral controls derived from three behavioral characteristics (speed, acceleration, and absolute angular velocity) and two different methodological approaches (mean differences shown in orange and means displayed in yellow). Rows 1-3 describe mean differences (method 1). For each animal, we calculated the average value of a behavioral variable (speed, acceleration, absolute angular velocity) for the sessions of interest. Subsequently, we calculated the absolute difference between the mean values for the sessions of interest and averaged to obtain a single value for a given animal. Rows 4-6 (method 2) describe average behavioral parameters. For each animal, we calculated the mean of the behavioral parameter of interest for the listed sessions. Subsequently, we averaged to obtain a single value describing the behavior of a given animal across the listed sessions.

**Figure 1:**
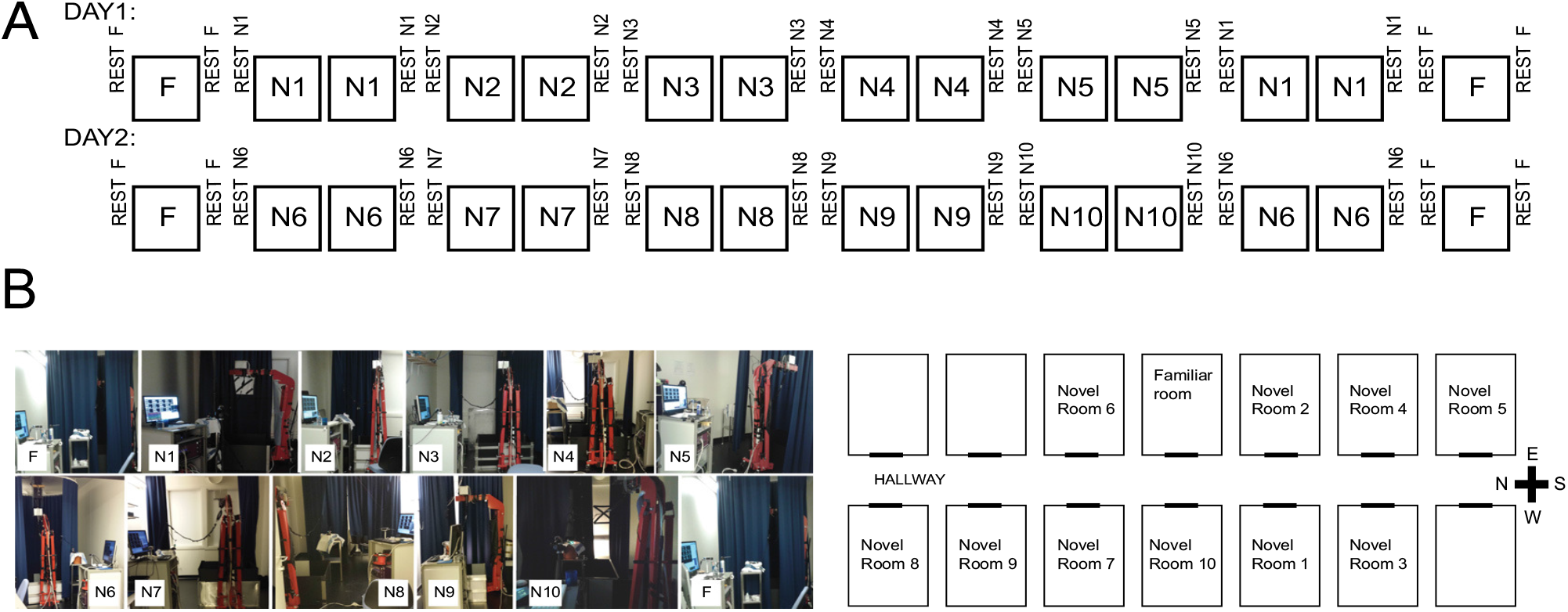
Experimental design. A) Experimental protocol. Five male Long Evans rats were exposed to 11 rooms over two days. In each room, the rats foraged for food in a 100 cm x 100 cm box. Animals were pre-trained in a familiar room (F) but did not experience any of the ten novel rooms (N) before testing. Exposure to each novel room was performed over two consecutive 15-minute sessions. The animal was not removed from the box between the first and second session. The familiar room (F) and the first novel room of each day, N1 and N6, were tested again at the end of the day. Each rest period lasted 15 minutes. Bl Pictures of the familiar and novel rooms. A mobile recording rig along with the red crane allowed continuous recording. The arrangement of the rooms along the hallway did not follow a particular pattern. Adapted from Alme et al. (2014).

**Figure 2:**
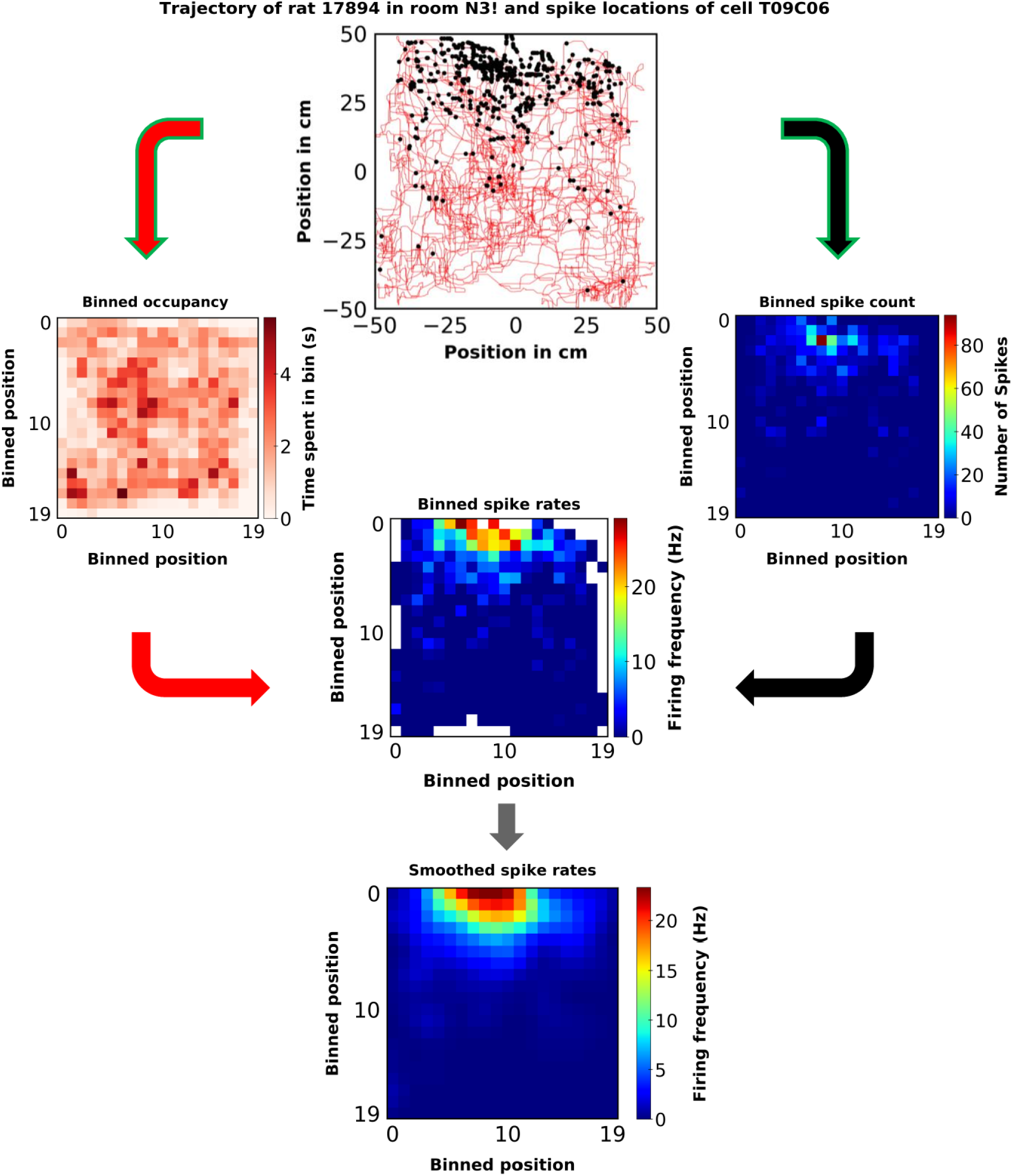
Data processing pipeline. The box in each session was binned using 400 (20 x 20) 5 cm square bins. Using the animals’ position data, we calculated each session’s binned occupancy in seconds (red arrows). Additionally, we counted how many times each place cell spiked in each bin (black arrows). The position data and the spiking data were speed-filtered (green line around the arrows). Periods during which the animals’ speed dropped below 5 cm/s were excluded from the binned occupancy, and spikes during these periods were not counted. Binned spiking rates for each place cell were calculated as the ratio of the number of spikes and the time spent in that bin in seconds (second red and black arrows). A Gaussian filter (sigma = 1 bin) was used to smoothe the rate maps (grey arrow).

**Figure 3:**
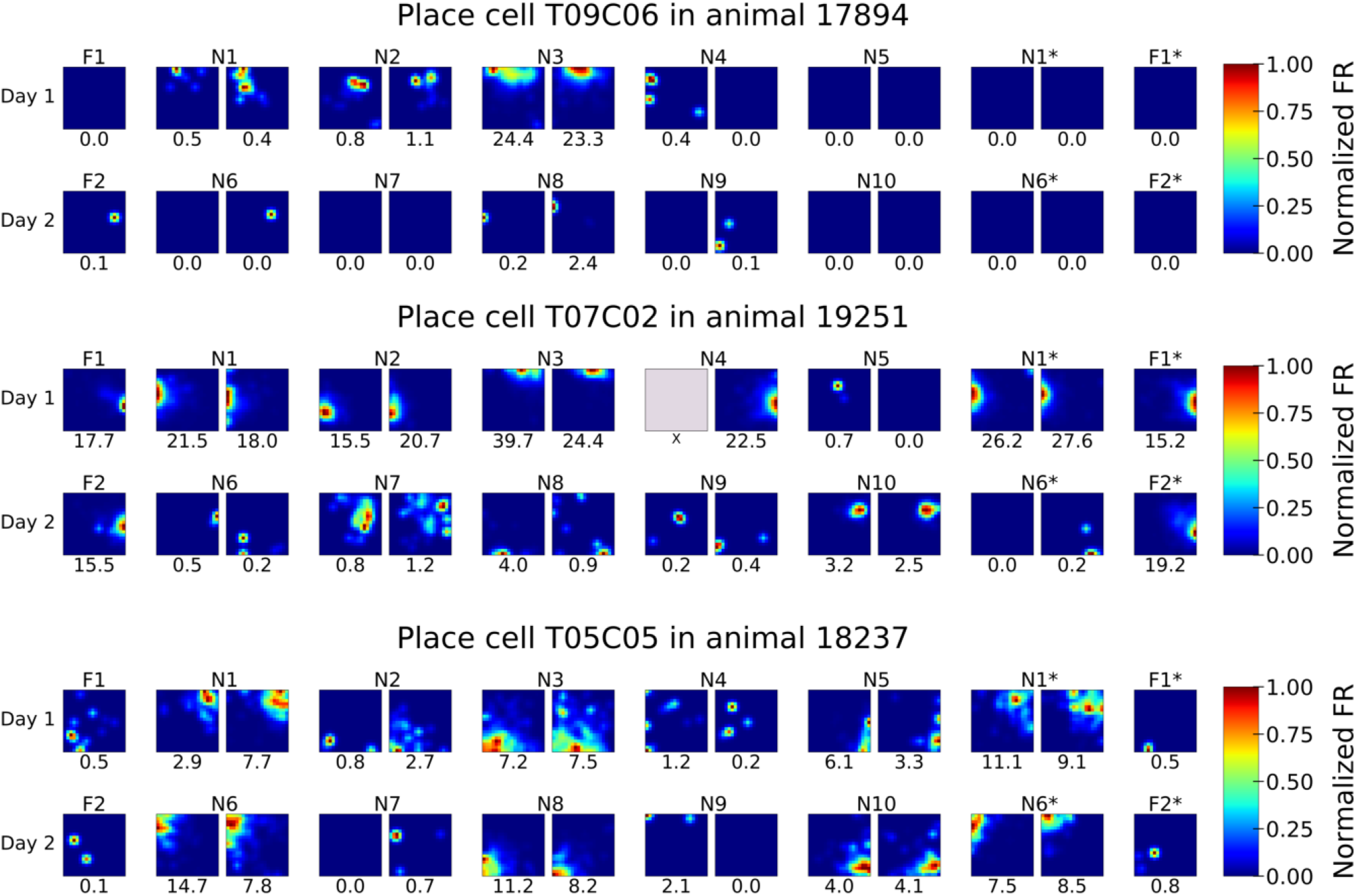
Smoothed firing rates of example place cells. We observed various instances of global and rate remapping. An instance of global remapping is the change of the place field location of place cell T07C02 in animal 19251 between the second session of N2 vs. the first session of N3 (RMC = −0.14). The same cell did not remap between the first and second session of N3 (RMC = 0.94). The remapping behavior of place cell T05C05 in animal 18237 between the first session of N1 and the third session of N1 later during the day is an example of rate remapping (RMC = 0.69). The color scale in each session is scaled to the maximum firing frequency displayed below the rate map. Grey boxes indicate unavailable data. F1, F1*, F2, and F2* all refer to the same familiar room.

**Figure 4:**
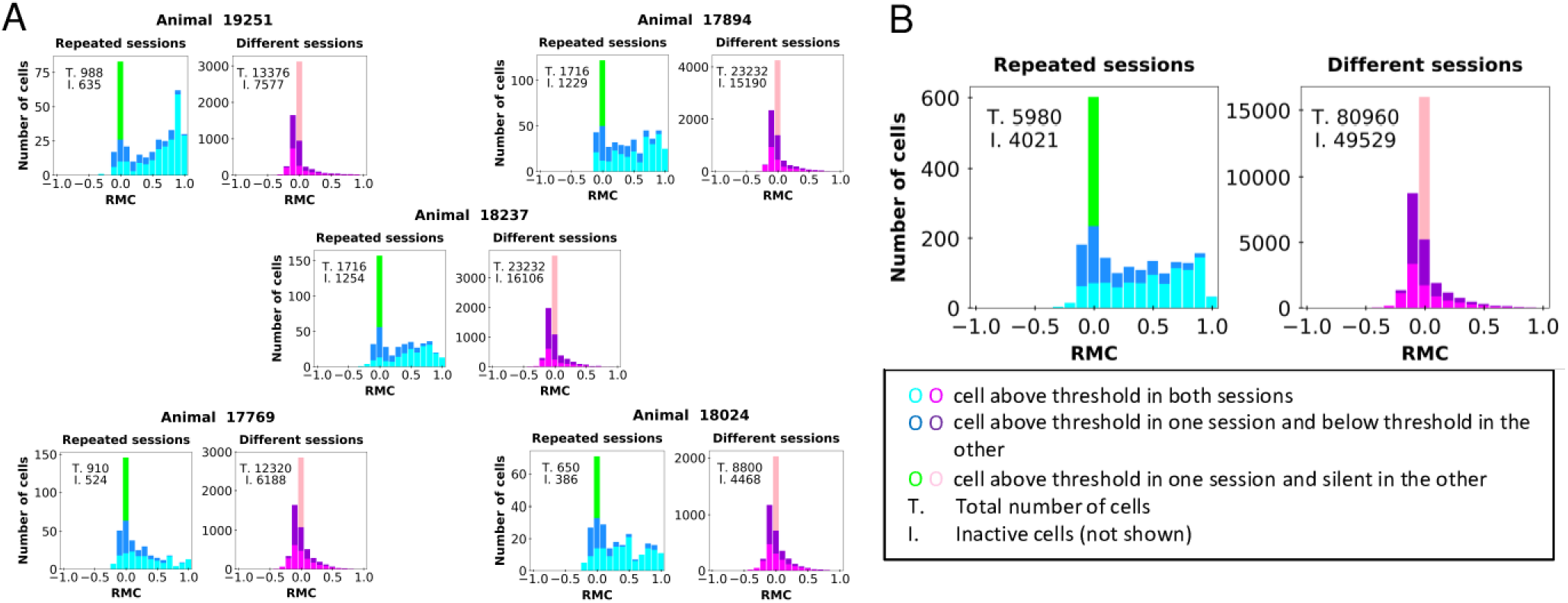
Rate Map Correlations (RMCs) across repeated and different sessions. Repeated sessions refer to repetitions of F, repetitions of each novel room, and repetitions of N1 and N6 at the end of each day. Different sessions include comparisons across different rooms. Total = total number of observations (number of recorded cells times the number of sessions compared). Inact. = inactive cells. Observations where the cell was either below threshold in both sessions, silent in both sessions, or silent in one and below threshold in the other session, were excluded from the analysis. A) RMCs of individual cells for each animal. In repeated sessions, RMCs of individual cells ranged from 0 to 1, indicating that partial remapping has occurred across the population of cells. This trend was consistent across all five animals. In different sessions, RMCs were clustered around 0, indicating that most cells remap. B) RMCs of individual cells across all animals. When pooling the data of all five animals, the pattern described in A) became more apparent.

2008), i.e., RMCs centered around 0.8-1. Yet, RMCs were relatively evenly distributed between 0 and 1 (Fig. 4). These findings support the notion that partial remapping occurs across repeated sessions. While some place cells remap (low RMCs), potentially indicating a new context in the same environment, other place cells do not remap (high RMCs), potentially coding for the unchanging spatial information. Interestingly, out of the cells that remap in repeated sessions, a large number of cells lose their place field entirely or gain a new place field (lime green bars in Fig. 4).

In order to compare different session pairs to each other, we generated a single number that described similarity in hippocampal maps across the session pair by averaging the RMCs across the population (Fig. 5). High average RMCs between two sessions indicate greater similarity in the hippocampal maps of these sessions and thus less remapping, whereas low average RMCs indicate greater dissimilarity and more remapping. Therefore, we expected to observe low average RMCs for comparisons of different rooms and relatively high average RMCs for repetitions of the familiar room (purple square near midline), immediate repetition of the novel rooms (turquoise squares one off diagonal), and repetitions of N1 and N6 at the end of each day (green four squares near midline). Although this pattern was broadly confirmed (Fig. 5), we observed variability among the five animals. Across 4 of the 5 animals, repetitions of F had high average RMCs, yet the magnitude of the correlation coefficients and thus the extent of remapping varied across animals. Immediate repetitions of the novel rooms showed consistently high average RMCs across all 5 animals. Yet again the remapping pattern was distinct for each animal, as indicated by the difference in the magnitude of the average RMCs across immediate repetitions of the novel rooms. Likewise, repetitions of N1 and N6 at the end of each day resulted in relatively high average RMCs in 4 of the 5 animals. Overall, the comparisons between different rooms led to low average RMCs or slightly negative average RMCs indicating that remapping occurs between different rooms (rel. white background in Fig 5). The exact remapping pattern was, however, variable and unique for each animal, as reflected by the uniqueness of each of the matrices. For some animals, we observed clusters of relatively high average RMCs, for rooms that were tested close in time to each other (light red boxes clustered together). This may be an indication that less remapping occurs between rooms that are tested closer to each other yet more data is needed to validate this claim.

**Figure 5:**
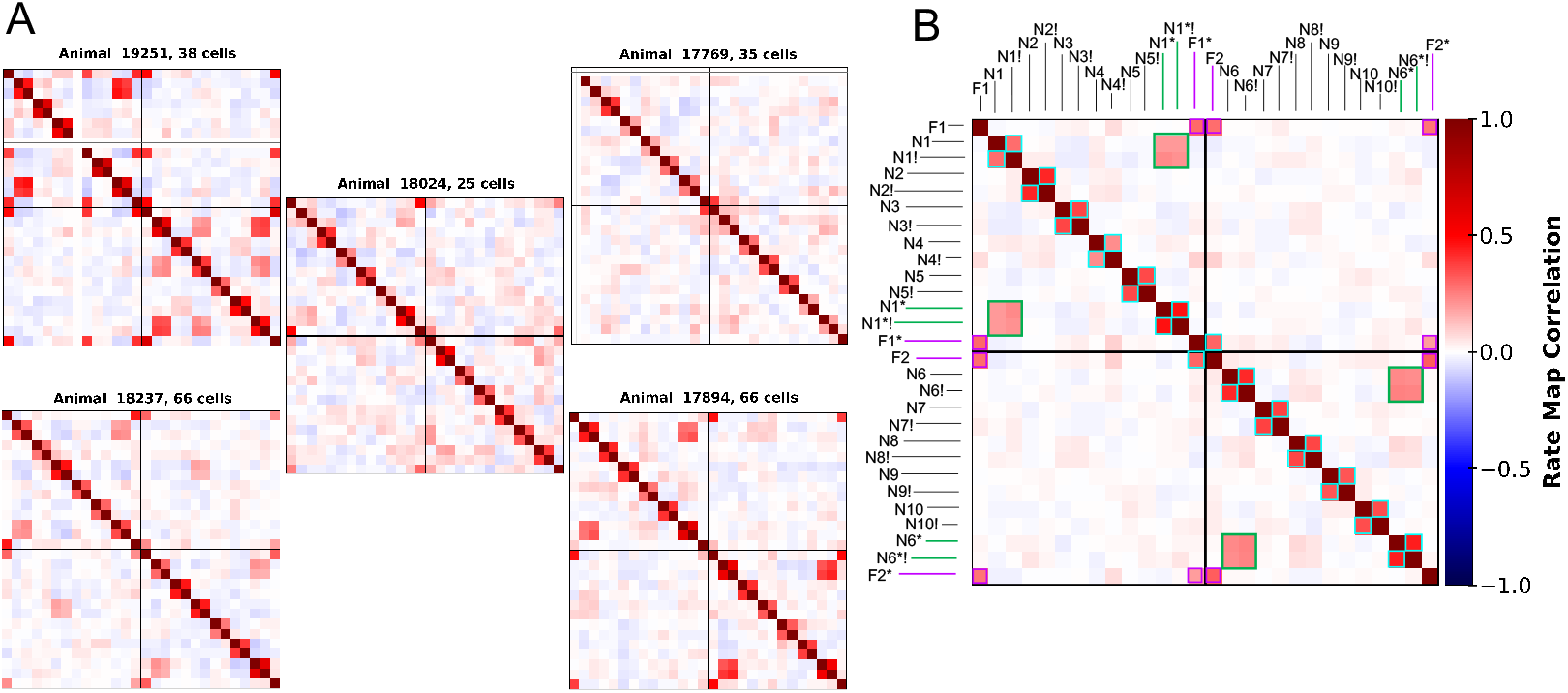
Average RMCs across animals and sessions. This is the population version of Fig. 4, illustrating the extent of remapping as a whole. A) Average RMCs of each animal individually. Overall, average RMCs were elevated for repetitions of the familiar room, immediate repetitions of the novel rooms, and repetitions of N1 and N6 at the end of each day. Comparisons between different sessions generally showed low average RMCs around 0. However, we observed a remarkable amount of deviation from this pattern across animals. In some animals, the repetitions of N1 and N6 at the end of each day displayed very low average RMC (animal 17769 and 18024). Additionally, in animal 19251 some comparisons between different rooms revealed relatively high average RMCs (see rooms N6 and N8). These findings indicate that remapping behavior is variable across animals. B) Average RMCs across all animals. When averaging mean RMCs across animals, the broad pattern described in A) became more apparent. While repetitions of novel rooms and the familiar room showed elevated average RMCs, comparisons between different rooms led to relatively low average RMCs. The repetitions of N1 and N6 at the ends of each day are marked with green boxes, whereas repetitions of the familiar room are marked with a purple frame. Immediate repetitions of the novel rooms adjacent to the diagonal are marked turquoise.

To examine the variability in animals’ remapping behavior, we monitored the distribution of average RMCs for repetitions of F vs. repetitions of N1 and N6 as described in Table 2 (Fig 6). We observed a very large range of average RMCs for both, repetitions of the familiar room (range ≈ 0-0.64) and repetitions of N1 and N6 at the ends of each day (range ≈ 0-0.47). Overall, we found that the mean RMCs for repetitions of F was higher than the mean RMC for the repetitions of N1 and N6 meaning that less remapping occurs in the familiar room. At the same time, we observed more variability (indicated by a wider distribution) in remapping behavior in the familiar room compared to the novel rooms N1 and N6. Both these findings were borderline significant and varied depending on the arbitrary changes in the parameters of our data analysis pipeline (e.g. speed-filtering and smoothing). More data is needed to make definitive conclusions about the animals’ remapping behavior in the familiar vs. the novel rooms. Interestingly, for the average RMCs of repetitions of N1 and N6 the animals’ average RMCs were clustered, raising the question if there is a pattern in animals remapping behavior.

**Figure 6:**
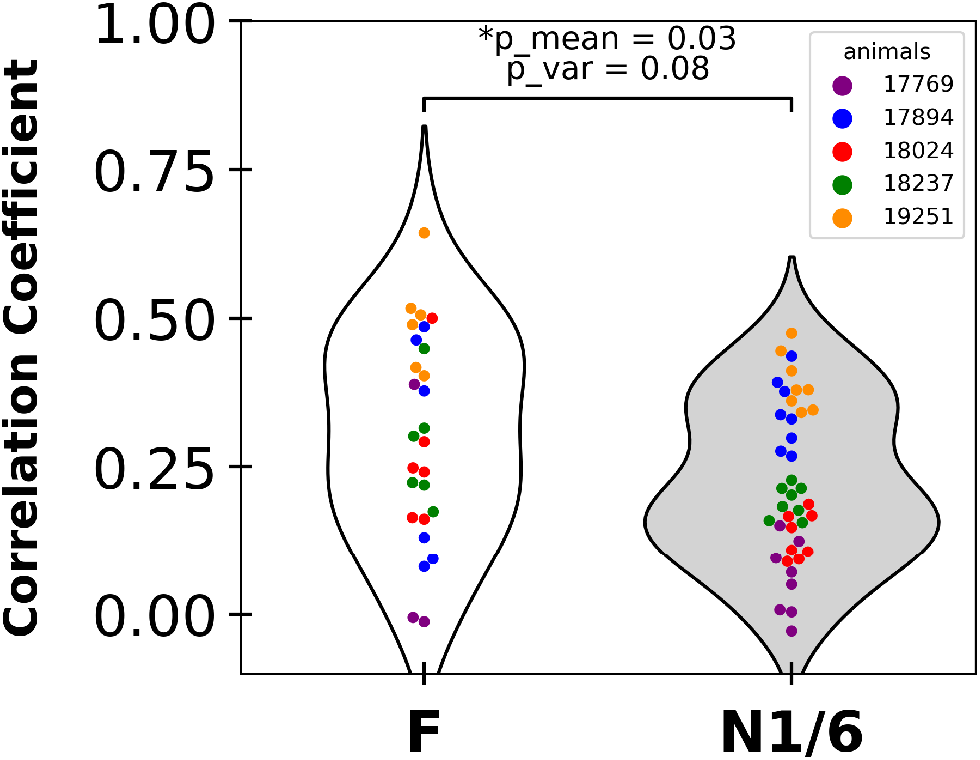
Remapping differences between familiar and novel rooms. The mean RMC for repetitions of F was higher compared to the mean RMC for the repetitions of N1 and N6 at the ends of each day. Furthermore, we observed more variability for RMCs describing repetitions of the familiar room relative to repetitions of N1 and N6 at the ends of each day. These findings were borderline significant and shifted based on arbitrary decisions concerning the speed-filtering, smoothing, and thresholding of the rate maps.

To examine the characteristics of remapping behavior across the population of animals, we looked at the consistency of RMCs across different comparisons in each animal. We found that animals with a higher mean RMC for the repetitions of F displayed higher RMCs for the repetitions of N1 and N6 at the ends of each day (Fig. 7A, *R*^2^ = 0.6, *p* = 0.08). Likewise, animals with a higher mean RMCs for repetitions of N1 showed higher RMCs for the repetitions of N6 (Fig. 7B, *R*^2^ = 0.72,*p* = 0.04). Additionally, animals with a higher Mean N within session RMC, showed a higher Mean N between session RMC (Fig. 7C, *R*^2^ = 0.93,*p* = 0.005). Taken together, these findings indicate that the remapping behavior across animals is structured and not random. Despite only 5 available data points (for 5 animals) in these analyses, the correlations described above were relatively high and stable across a variety of values of the parameters of our pre-processing pipeline (e.g., speed-filtering, smoothing).

The number of recorded cells varied across the five animals (Table 1; range: 25-66 cells). To test whether the size of the recorded place cell population affected animal-to-animal variability, we performed a random sampling analyses (for detailed procedures see materials and methods section). We took 100 random samples of 25 cells from one of the animals with the highest number of cells and calculated the average RMC for F1/F2 for each random sample (Fig 8A). We compared that distribution of subsampled population average RMCs to the average RMC for F1/F2 calculated with the full population for that animal and for the animal with the smallest population (18024, 25 cells). If variation in recorded population size caused the variation in RMCs, the average F1/F2 RMC of the animal with the lowest number of cells (18024, red line) should have fallen within the distribution of the average RMCs of the animal with the highest number of cells after subsampling to the same population size (17894, blue bars). However, the average F1/F2 RMC of animal 18024 was clearly outside the range of the subsampled population RMCs, whereas the average F1/F2 RMC of animal 17894 was in the center of the range of the subsampled population RMCs, indicating that variation between animals in the number of cells sampled did not give rise to the variation between animals in remapping behavior. Furthermore, we examined whether sampling bias affected the study’s main finding, shown in figure 7. Because the animal with the lowest number of cells had 25 recorded units, we reconstructed the correlation shown in Fig. 7C 100 times, each time sampling 25 random cells from the other four animals. The subsampled data of each animal was clearly clustered around the original data point and the variability across animals was preserved (Fig. 8B). Taken together, these findings suggest that variation across animals in recorded population size is unlikely to explain the structured variability in the animals’ remapping behavior.

**Figure 7:**
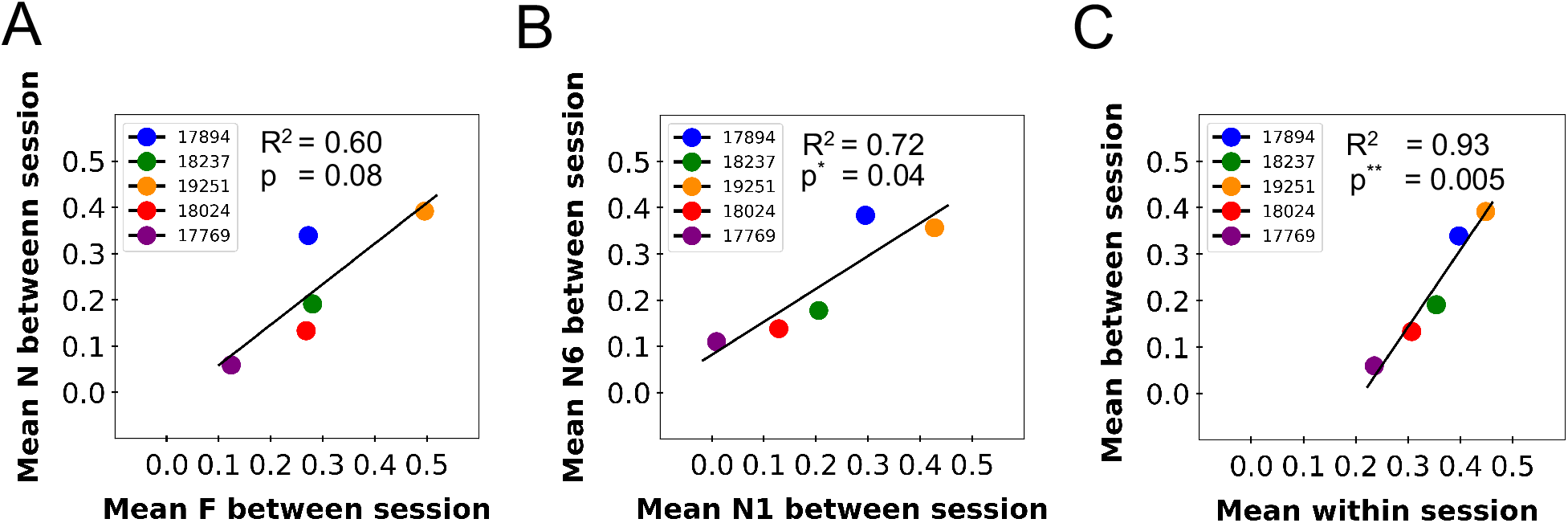
Structured Heterogeneity in animals’ remapping behavior. A) Relationship between mean F and mean between session RMC. Rats with higher RMCs for repetitions of F displayed higher RMCs in the repetitions of N1 and N6 at the end of each day. B) Relationship between mean N1 and mean N6 RMCs. Rats with higher RMCs for repetitions of N1 at the end of day 1 displayed higher RMCs for N6 at the end of day 2. C) Relationship between mean within session and mean between session RMCs. Animals with higher within session RMC (immediate repetitions) displayed a higher between session RMC for repetitions of N1 and N6 at the end of each day.

To test whether differences in behavior accounted for the differences in remapping shown in Figures 5-7, we quantified animals’ average speed, acceleration, and absolute angular velocity for each session. We wanted to test whether the correlations between the different RMCs for each animal were actually due to correlations of those RMCs or due to some other behavioral variable that was consistent in each animal. To do so, we constructed Generalized Linear Models (GLMs) that augmented each of the correlations shown in Figure 7C with one of these behavioral variables. For each GLM, there were two independent variables (regressors) and one dependent variable. Exactly like the correlations in Figure 7, the dependent variable was the average RMCs for a particular set of session pairs averaged across all pairs of that class. Like the correlations in Figure 7, one of the independent variables was the average RMCs for a different set of session pairs averaged across all pairs of that class. In addition, the other independent variable was the difference in one of the behavioral variables between sessions in the same session pairs as those used for the dependent variable, averaged across all session pairs in that class. The idea behind structuring the GLMs this way is that we wanted to see whether the variability in the neural data for a given set of session pairs was better predicted by behavioral variability in that same set of session pairs or by variability in the neural data for a completely different set of session pairs. We used two classes of behavioral control variables, constructed as explained in Table 3. One class was the mean differences in the behavioral variables (Table 3, orange). The mean differences in a behavioral variable between two sessions reflect the dissimilarity with respect to that behavioral variable, just as RMCs reflect the similarity in hippocampal maps between two rooms. Therefore, using mean differences as a control for RMCs compares the similarity of behavior across two sessions with the similarity of hippocampal maps between those same sessions. The other class was the mean behavioral variables themselves (Table 3, yellow), which allow us to describe the animal’s average behavior across all the sessions that were used to construct the neural variables used for the GLMs in figure 7C. In this approach, we controlled for the possibility that animals’ behavioral characteristics may directly relate to RMC values, as would be expected for example if average speed and RMCs both related to attentional levels. In each of the six controls of 7C, we observed a high correlation between the neural independent variable and the neural dependent variable along with a low correlation between the behavioral independent variable and the neural dependent variable (Table 4). This finding confirmed that the high correlation between Mean N within session RMCs and Mean N between session RMCs cannot be explained by the animals’ speed, acceleration, and absolute angular velocities, nor can it be explained by changes in those behavioral variables. The exact structure of the GLMs can be found in Table 3.

**Figure 8:**
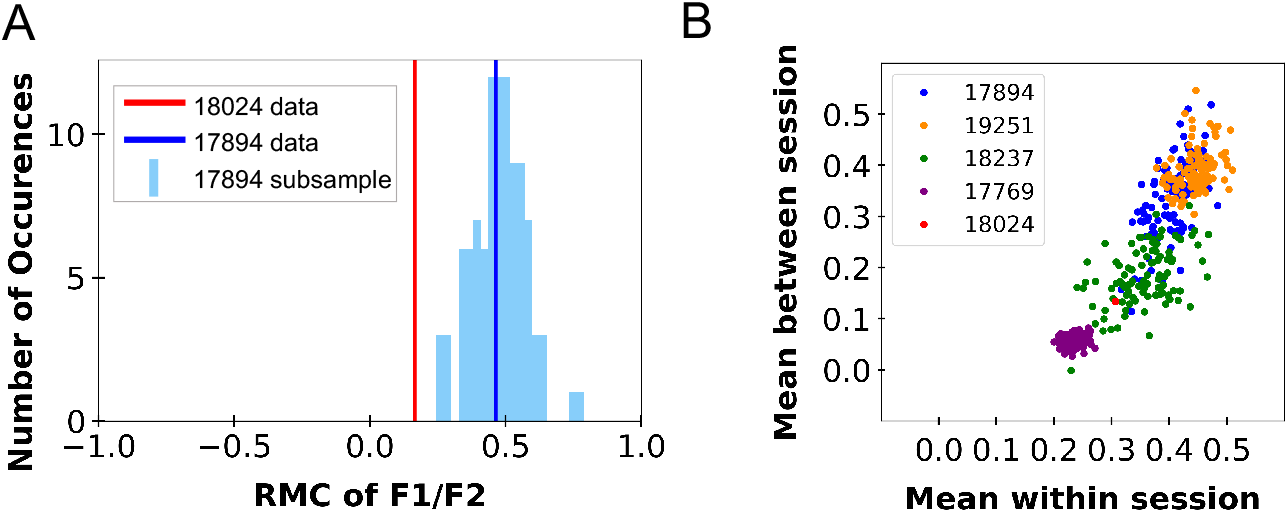
Experimental Sampling Controls. A) We compared the animal with the fewest recorded cells (18024, 25 cells) with 100 random subsamples of 25 cells from one of the animals with the most cells (17894, 66 cells). The bars show the number of occurrences of each average F1/F2 RMC among the 100 subsamples of animal 17894 (blue). The vertical lines are the average RMC for the F1/F2 session pair for the full recorded population for each animal (17894 - Blue, 18024 - red). The full population RMC for animal 17894 is at the center of its own subsampled range of RMCs, whereas the full population RMC for animal 18024 is outside the range of subsampled RMCs for animal 17894. B) We recreated a subsampled version of Fig. 7C where each dot is the RMC calculated using a subsample of 25 cells from that animal. As evident by the linear arrangement of the data, higher mean-within session RMCs correlated with higher mean-between session RMCs, and the data for each animal maintained the order of Fig. 7C.

## Discussion

Hippocampal remapping is a complex and highly variable phenomenon. In this work, we studied the characteristics of cell-to-cell and animal-to-animal variability in remapping behavior. We compared the firing patterns of individual cells of five rats across 11 repeated and different rooms. While most cells remapped across different room comparisons, we observed a large amount of variability across repeated rooms, indicating that partial remapping occurs in the latter type of comparison. We then shifted our focus from individual cells to comparing the changes in the hippocampal maps of the animals across repeated and different sessions, using average RMCs. We discovered extensive remapping across different rooms, but much less remapping across repeating sessions, indicating that hippocampal maps are similar yet not identical across repeated sessions. Although this general remapping pattern applied to all five animals, we detected variability across the animals. We quantified the pattern of variability in the animals’ remapping behavior by correlating different categories of neural remapping data (Fig. 7). Across all comparisons, we discovered a high correlation between the neural variables of the five animals, and the animals stratified along a spectrum that was preserved for each comparison. We concluded that the animal-to-animal variability in remapping behavior is structured. Neither the subsampling of place cells (Fig. 8) nor the animals’ underlying behavioral characteristics (Table 4) accounted for the structured variability we observed, raising the question of the origins of this phenomenon.

**Table 4:** Variables used in GLMs of Figure 7. Each row describes how we calculated the value of a variable used in Figure 7. The rows contain the name of the variable in the left column, a verbal description of the session pairs used in the middle column, and the actual list of session pairs in the right column. For a given variable, we calculated the RMC for each cell for the session pairs listed in the right column. We averaged across all cells for each session pair. We then averaged across all session pairs for each animal. In this way, we arrive at a single value of each variable for each animal.

| Ind. Var. 1 | Ind. Var. 2 | Dep. Var. | R<sup>2</sup> (1) | P_val (1) | m (1) | P_val (2) | m (2) |
| --- | --- | --- | --- | --- | --- | --- | --- |
| Mean N within session | N/A | Mean N between session | 0.93 | 0.005 | 1.66 | N/A | N/A |
| Mean N within session | Mean speed difference N between session | Mean N between session | 0.97 | 0.04 | 1.22 | 0.15 | -0.12 |
| Mean N within session | Mean acceleration difference N between session | Mean N between session | 0.95 | 0.05 | 1.32 | 0.30 | -0.06 |
| Mean N within session | Mean absolute angular velocity difference N between session | Mean N between session | 0.91 | 0.04 | 1.76 | 0.75 | 0.0013 |
| Mean N within session | Mean speed N between session | Mean N between session | 0.92 | 0.05 | 1.49 | 0.58 | 0.01 |
| Mean N within session | Mean acceleration N between session | Mean N between session | 0.93 | 0.05 | 1.47 | 0.50 | 0.01 |
| Mean N within session | Mean absolute angular velocity N between session | Mean N between session | 0.94 | 0.02 | 1.66 | 0.35 | -0.0018 |

### Possible origins of animal-specific remapping behavior

There are several potential explanations for this animal-specific remapping behavior that we cannot yet distinguish between.

One class of possibilities (“strategy variability”) is that animals have computational-level differences in how they deal with the inherent ambiguity in context definition. As described in the Introduction, animals do not have direct access to context labels, and therefore must *infer* the hidden context identity, a process characterized as hidden state inference by Sanders et al. (2020). The tradeoff between “splitting” and “lumping” (increased differentiation vs increased generalization, more vs fewer hidden states) has no a priori solution, which is why populations will have individuals with a variety of tendencies on that axis (Simpson, 1945). Sanders et al. (2020) suggested that differing settings of a parameter of the model, alpha, could underlie variation in remapping behavior as it controls the tendency of the model to prefer a greater or smaller number of hidden states. Of course, other forms of variation could arise from other parameters of that model or of other models of remapping behavior that explicitly include parameters with no a priori optimal values. In short, this class of possibilities suggests that variability in remapping behavior might correspond to variability in problem-solving strategies.

Another class of possibilities (“capability variability”) is that animals have differences in cognitive abilities. Learning impairments can cause an animal to be unable to retrieve a memory of an environment. If that occurs, the map would not be reused when the animal re-enters the environment, which would result in remapping (near-zero average RMC). A previous result interpreted using this perspective is a series of studies on aging rats by Barnes et al. (1997, reviewed by Lister and Barnes, 2009), who show that aging animals have a greater tendency to remap when presented with the same environment. Indeed, others showed that among aged animals, tendency to remap correlated with other firing characteristics (Hok et al., 2012). In this way, it could be that the variability across animals observed in our data was the result of learning deficits in some animals. A related explanation in this class is that remapping can be induced by a lack of attention paid to the cues that differentiate environments. For example, Kentros et al. (2004) show that changing attentional demands changed the tendency of animals to remap. The experimental data we analyzed were recorded during a task with low attentional demands, so intrinsic differences in attention between animals may have given rise to differences in remapping tendencies. This class of possibilities suggests that variability in remapping behavior might correspond to variability in cognitive capabilities.

The key distinction between these classes of hypotheses is how variability in neural responses corresponds to variability in behavior. The “strategy variability” hypothesis would suggest that different animals would be differentially skilled at different tasks. The “capability variability” hypothesis would suggest that some animals would be better at all context-dependent tasks.

Unfortunately, it is unclear exactly how remapping relates to behavior. The standard assumption in the field is that hippocampal remapping corresponds to context-specific learning: that animals will generalize behaviors learned between experiences using the same map but not between experiences using different maps (Colgin et al., 2008). This assumption has not been directly proven, and in fact there is limited evidence to the contrary (Jeffery et al., 2003). For the sake of argument, let’s accept this assumption, however. Under the “capability variability” hypothesis, animals who remap more would simply have more difficulty generalizing knowledge past the experience they learned it in, as they do not reuse maps. Under the “strategy variability” hypothesis, something more nuanced occurs. Animals with a greater tendency to remap would be faster at learning tasks that required distinctions, such as a context-specific go-no-go task. This would be because animals that don’t remap would have a greater tendency to generalize across slightly different experiences. On the other hand, animals with a lower tendency to remap would be faster at learning tasks that required greater levels of generalization. Another potential behavioral prediction of the “strategy variability” hypothesis is that the tendency of remapping might correspond to the extent of generalization during fear conditioning. When a shock is only presented a single time, the animal has to infer what extent of generalization of that experience is appropriate. Differences in sensitivity in environmental changes giving rise to remapping would be predicted to correspond to differences in sensitivity to environmental changes that elicit a conditioned fear response.

One final possibility is that differences in the recording locations in different animals may have given rise to the differences in remapping behavior, if different parts of the hippocampus respond characteristically differently. There are reasons to believe that remapping might have different properties along the CA3-CA1 (proximo-distal) axis Lee et al. (2004); Leutgeb et al. (2004); Guzowski et al. (2004) as well as along the dorsal-ventral (septo-temporal) axis Royer et al. (2010). If these different animals were implanted in characteristically different locations, the remapping behavior of the recorded cells might be consistently different even if the animals in general did not have consistent differences. We do not have access to the recording locations of the animals, so we could not verify this hypothesis.

One result that gives rise to several questions is that of the subsampling done in Fig. 8. In addition to the across-animal correlation that is preserved despite subsampling, there is also within-animal correlation across separate subsamples. This implies that different cells within an animal have different tendencies to remap. This could arise with the previous hypothesis, that different recording locations might have characteristically different remapping behaviors. A related possibility was mentioned in Sanders et al. (2020), namely, that hierarchical hidden state inference could be performed by having a gradient of remapping tendencies within a population.

Finally, this paper suggests a “poor man’s” remapping metric: within-session variability. We show that within-session remapping correlates with between-session remapping (Fig. 7C). If between-session remapping tendency turns out to correlate with learning style (as suggested by the “strategy variability” hypothesis) or with learning capability (as suggested by the “capability variability” hypothesis), measures of within-session firing variability such as overdispersion (Olypher et al., 2002; Kelemen and Fenton, 2016) could be used as a proxy for tendency to notice small differences or with learning speed, respectively. Indeed, remapping has been shown to correlate with overdispersion in aged rats (Hok et al., 2012). The utility of this measure awaits research into its behavioral relevance, but certainly future studies of remapping should keep it in mind.

## Acknowledgments

This material is based upon work supported by the Center for Brains, Minds and Machines (CBMM), funded by NSF STC award CCF-1231216.

